# Engineered ADARs enable single-nucleotide resolution DNA A-to-G editing without bystander effects

**DOI:** 10.1101/2025.07.19.664718

**Authors:** Hyeon Woo Im, Bada Jeong, Yeji Lee, Ye Eun Oh, Chanju Jung, Yong-Woo Kim, Heesoo Uhm, Sangsu Bae

## Abstract

The adenine base editor (ABE), which enables A•T-to-G•C base conversion, has emerged as a powerful tool for therapeutic applications. However, conventional ABEs suffer from bystander nucleotide conversions, limiting their utility for precise editing. Here, we present a single-nucleotide resolution ABE (snuABE) created by fusing a nickase Cas9, nCas9(H840A), with the deaminase domain of ADAR, which acts on DNA:RNA hybrids, instead of TadA, which acts on single-stranded DNA in conventional ABEs. snuABE requires a specially designed target-adenine guide RNA (tagRNA) that introduces a mismatch at the target adenine, enabling highly specific A-to-G editing by ADAR. Engineering ADAR from Pediculus humanus using the in silico protein evolution algorithm EvolvePro, along with 3’-end protection of the tagRNA, further enhances the editing activity of snuABE in human cells. Moreover, snuABE exhibits significantly reduced DNA off-target activity, highlighting its potential as a safer and more precise base editing technology for therapeutic applications.

## Main

The CRISPR-Cas system has been widely used for genetic engineering with high efficacy. CRISPR nucleases are designed to introduce DNA double-strand breaks (DSBs) at target sites in a guide RNA (gRNA)-dependent manner, primarily resulting in gene disruption^1^. However, this strategy is not suitable for many genetic diseases that require gene correction or base conversion. Additionally, CRISPR nuclease-induced DSBs can lead to serious safety concerns, including large DNA deletions^2,3^, chromosomal aberrations^4^, and p53-mediated cell death^5^.

As an alternative, base editors (BEs) have gained significant attention because they enable the modification of one or a few nucleotides with high editing efficiency, without inducing DNA DSBs^6^. BEs typically consist of a nickase Cas9 (nCas9)(D10A) fused to a specific deaminase that catalyzes nucleotide conversions. However, BEs still face notable limitations, which have been partially addressed through several strategies: (i) gRNA-dependent off-target editing in the genome ^7^, which can be reduced by employing high-fidelity Cas proteins and modified gRNAs^8^; (ii) gRNA-independent off-target editing in the genome^9,10^ and transcriptome^11–13^, which can be mitigated using engineered deaminases^13^; and (iii) unintended base conversions near the target site (bystander editing) within the editing window, which can be minimized using BE variants with narrower activity windows^14^. However, despite substantial efforts, completely eliminating bystander editing remains a challenge and often hampers functional recovery^15^. Furthermore, a trade-off exists between editing efficiency and specificity: BEs with enhanced editing activity typically show increased bystander editing. These limitations underscore the need for the development of a highly precise base editing tool that can achieve accurate gene correction without bystander effects.

ADARs (adenosine deaminases acting on RNA) selectively recognize and deaminate mismatched adenosines to inosines (A-to-I editing) in double-stranded RNA, a process that can alter protein function, affect RNA stability, and influence gene expression^16^. Leveraging this property, several RNA adenine base editing tools such as CIRTS, LEAPER, RESTORE, REPAIR, and RESCUE have been developed ^17–21^. However, to the best of our knowledge, ADAR has not yet been applied for DNA adenine base editing in mammalian cells. In this study, we introduce a single-nucleotide resolution adenine base editor, termed snuABE, in which the deaminase domain of ADAR is fused to the N-terminus of nCas9(H840A). Unlike conventional ABEs that use single-guide RNAs (sgRNAs), snuABE employs a specialized gRNA, target-adenine guide RNA (tagRNA), which creates a mismatch at the target adenine site. For a given DNA target, snuABE can selectively convert specific adenines by designing tagRNAs that introduce unique target-specific mismatches. Notably, the overall editing efficiency of snuABE was enhanced using EvolvePro^22^, an artificial intelligence (AI)-driven directed evolution framework. Furthermore, snuABE demonstrated extremely low levels of DNA off-target editing activity, highlighting its therapeutic potential as a precise and safe base editing tool.

## Results

### Operating concept of snuABE utilizing ADAR and a dedicated guide RNA

Although ADAR typically acts on double-stranded RNA (**Supplementary Fig. 1a**)^16^, a previous study has shown that it can deaminate DNA adenines within DNA:RNA duplexes *in vitro*^23^. Based on this finding, we hypothesized that ADAR could be repurposed for DNA adenine editing in human cells, provided that the target adenine is flipped out in an DNA:RNA hybrid conformation (**Fig. 1a**). To test this, we designed six constructs in which the deaminase domain of human ADAR2 (hADAR2dd) was fused to either the N- or C-terminus of various Cas9 variants, including nCas9(H840A), nCas9(D10A), and catalytically inactive Cas9 (dCas9) bearing both D10A and H840A mutations (**Fig. 1b**). To create a specific mismatch at the target adenine, we engineered a tagRNA by extending the 3’-end of a conventional sgRNA. We next evaluated the six snuABE constructs using a tagRNA designed to introduce a mismatch at the 12^th^ adenine of a FANCF target site in HEK293T cells. While ABE8e, the representative ABE varianta, edited multiple adenines at positions 12, 13, and 18 with efficiencies of up to 61.0%, snuABE exhibited lower but selective editing activity (1.1%) exclusively at the 12^th^ adenine. These results confirmed that snuABE can function in human cells with high positional specificity. This initial version was named snuABE1.0.

**Fig. 1.**
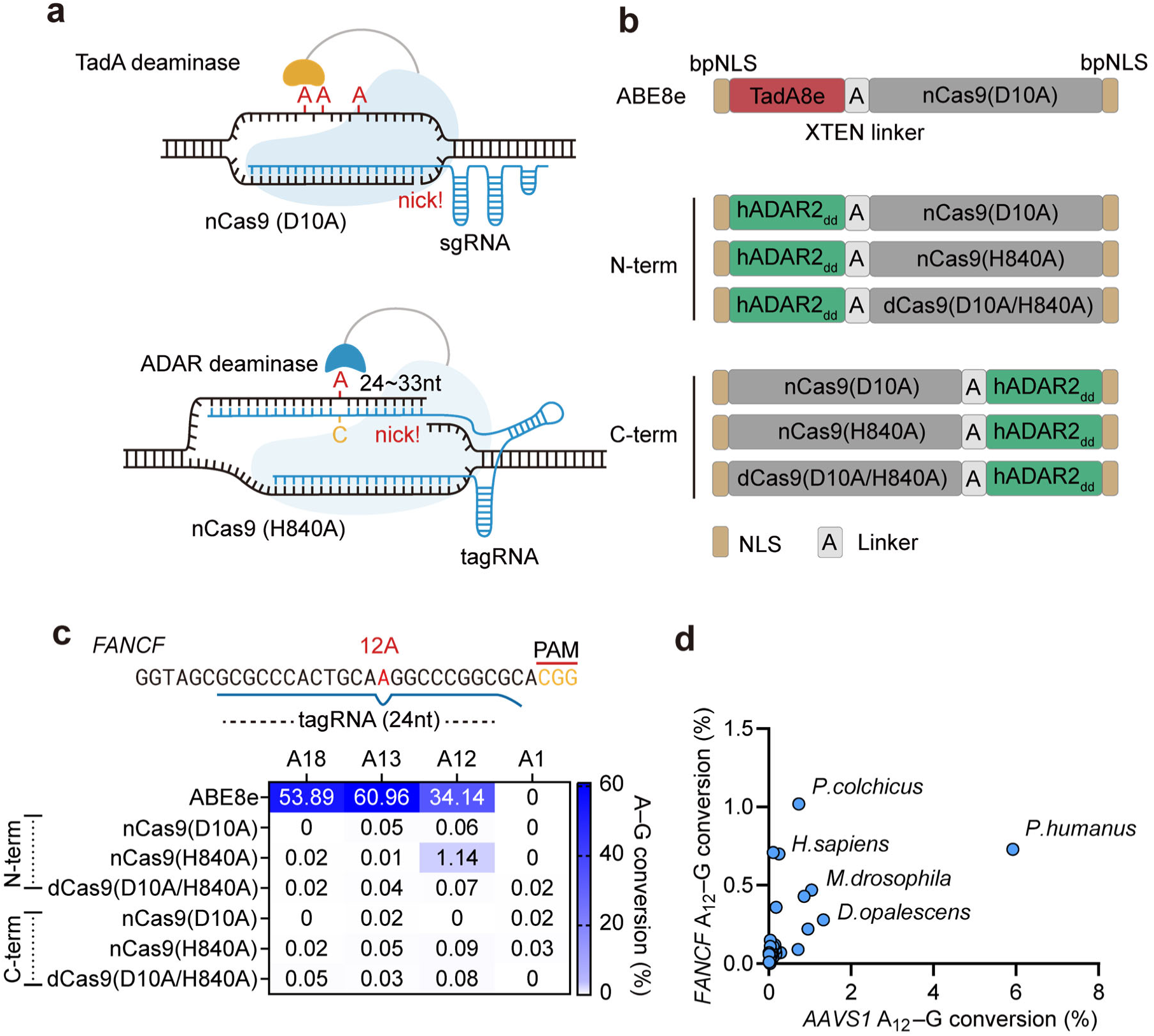
Conceptualization of a novel base editor (snuABE) utilizing ADAR and cross-species ADAR screening. **a,** Illustration comparing conventional ABEs and snuABEs, the latter utilizing an ADAR deaminase domain. **b**, Schematics of conventional ABE8e and snuABEs based on the human ADAR2 deaminase domain (ADAR2dd). **c**, Comparison of editing efficiencies at the human *FANCF* locus using the constructs shown in **b**. **d**, Editing efficiency comparison of different ADAR proteins tested at the human *AAVS1* and *FANCF* loci. Cell values in heat maps represent the mean, while bar graphs show the mean ± s.d. with error bars (*n* = 3 independent biological replicates). All experiments were performed in HEK293T cells.

ADAR enzymes were screened to identify variants with improved editing activity. Given that ADARs are found across a wide range of eukaryotes, 22 ADAR deaminase domain (ADARdd) orthologs from 20 different species, including primates, insects, and marine organisms, were retrieved from the NCBI database. Previous studies have reported that substituting a specific glutamic acid residue with glutamine (e.g., E488Q in hADAR2dd) enhances deaminase activity ^24^. Therefore, we introduced analogous E-to-Q substitutions at conserved residues across all 22 ADARdd orthologs **(Fig. 1d** and **Supplementary Fig. 1)**. Remarkably, among the tested ADARdds, the deaminase domain from *Pediculus humanus* (PhADARdd) demonstrated superior editing activity at both the FANCF target (0.7%) and the AAVS1 target (6.0%) **(Fig. 1d)**. This improved version was named snuABE2.0.

### 3’-End protection of tagRNA increase snuABE editing efficiency

Although snuABE demonstrated higher specificity than ABE8e, its relatively low editing efficiency prompted further optimization. To improve the performance of snuABE2.0, we adopted several strategies previously used to develop the optimized prime editing (PE) system, PEmax^25^. Specifically, we generated snuABE2.1 by incorporating the SV40 internal nuclear localization signal (NLS), introducing two point mutations (R221K and N394K) into nCas9(H840A), and adding a C-terminal c-Myc NLS (**Fig. 2a**). Given the structural similarity between tagRNA and the PE guide RNA (pegRNA), we hypothesized that protecting the 3’-end of tagRNA might further enhance editing efficiency^26^. To test this, we co-expressed full-length La or truncated La(1–194) protein along with snuABE2.1. At both the *FANCF* and *AAVS1* target sites, snuABE2.1 showed improved editing efficiency over snuABE2.0, and co-expression with either form of La protein further enhanced this activity (up to 8.6% for *AAVS1* and 5.1% for *FANCF*) (**Fig. 2b**, **2c**). Encouraged by these results, we next directly fused the truncated La(1–194) domain to various positions within snuABE2.1, including the N-terminus, internal linker region, and C-terminus (**Fig. 2a**). The highest editing efficiency was observed when La(1–194) was fused to the C-terminus, reaching 16.5% for *AAVS1* and 8.5% for *FANCF*, which was higher than with separate co-expression of La(1–194) (**Fig. 2b**, **2c**). This optimized version was designated snuABE3.0. Based on snuABE3.0, we developed snuABE3.1 by applying additional PEmax-inspired modifications. This final construct achieved robust editing efficiencies of ∼20.0% for *AAVS1* and 12.1% for *FANCF* (**Fig. 2b**, **2c**).

**Fig. 2.**
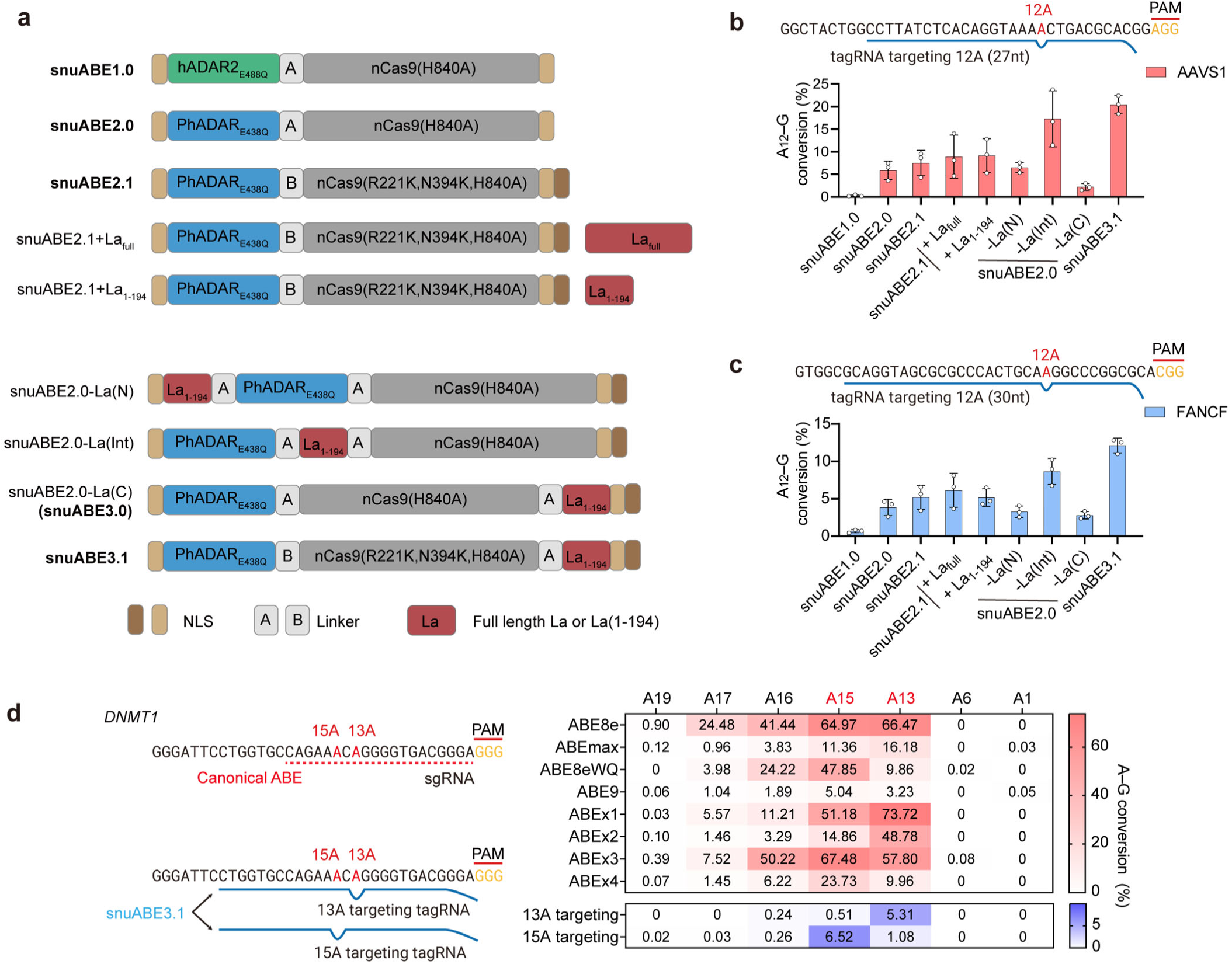
Engineering of snuABEs and target adenine selectivity through tagRNA switching. **a**, Engineering schemes of snuABE constructs from versions 1.0 to 3.1. **b**,**c**, Comparison of editing efficiencies at the human *AAVS1* and *FANCF* loci using constructs from **a**. **d**, Comparison of snuABEs with conventional ABEs at the endogenous *DNMT1* locus. ABEx1–x4 were used with agRNAs, other ABEs with sgRNAs, and snuABEs with tagRNAs. Cell values in heat maps represent the mean, and bar graphs display the mean ± s.d. with error bars (*n* = 3 independent biological replicates). All experiments were performed in HEK293T cells.

We next assessed whether snuABE could selectively edit individual adenines at a given target site by introducing position-specific mismatches using tailored tagRNAs. For this purpose, we designed two tagRNAs targeting either the 13^th^ or 15^th^ adenine at the same site in *DNMT1*. In parallel, we evaluated a panel of conventional ABEs, including ABEmax, ABE8e, ABE8eWQ, ABE9, and recently developed bystander-reduced variants ABEx1–4^14,27–30^. While all conventional ABEs and ABEx variants edited multiple adenines within their activity window, snuABE3.1 demonstrated clear single-nucleotide resolution. Specifically, with the tagRNA targeting the 13^th^ adenine, snuABE3.1 efficiently edited the 13^th^ adenine (5.3%) while sparing the 15^th^ (0.5 %). Conversely, the 15^th^ adenine-specific tagRNA yielded high editing at the 15^th^ position (6.5%), with minimal editing at the 13^th^ position (1.1 %) (**Fig. 2d**). These results strongly support the unique single-nucleotide selectivity of snuABE3.1.

### Characterization of optimized conditions of snuABE3.1

We next investigated various characteristics of snuABE3.1, including the optimal tagRNA length, base editing purity, editing window, and sequence motif preference. When testing tagRNAs with 3’ extensions ranging from 15 to 69 nucleotides (nt), the highest editing efficiencies were consistently observed with tagRNA lengths between 24 and 33 nt (**Fig. 3a** and **Supplementary Fig.2**). Similar to conventional ABE8e, snuABE3.1 induced minimal insertion and deletion (indel) events (<1.0%) (**Fig. 3b**), suggesting that snuABE3.1 has a low risk of causing genomic instability. Furthermore, A-to-G conversion was the predominant outcome at target sites, with minimal non-specific base conversions (**Fig. 3b**). To investigate base-pairing requirements for efficient editing, we tested various counter-RNA bases at the position opposite the target DNA adenine in the mismatch. Only cytosine led to efficient A-to-G conversion, while other bases contributed little to no editing activity (**Fig. 3c**), consistent with previous observations^19^. To identify the effective editing window, cells were treated with snuABE3.1 using 32 tagRNAs targeting adenines at different positions. The highest editing efficiencies were consistently observed when adenines were located between positions 11 and 16 **(Fig. 3d** and **Supplementary Fig. 3a–m)**. Notably, when adenines at positions 18 and 19 were targeted, unintended bystander editing occurred at adenines located within positions 12 and 14 (**Supplementary Fig. 3j–3i**), further supporting a confined editing window spanning positions 11 to 16.

**Fig. 3.**
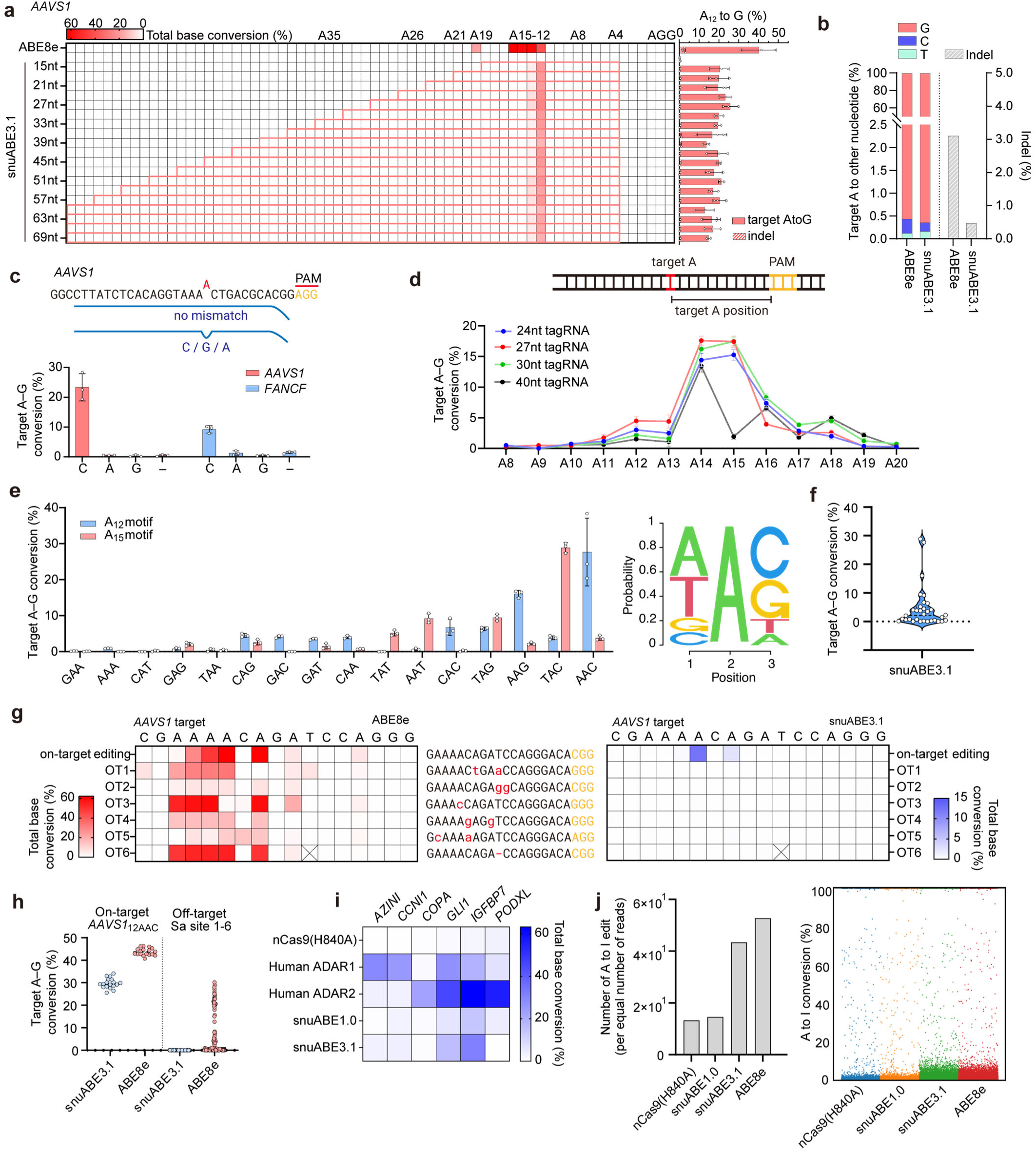
Functional characterization of snuABE in genomic and transcriptomic contexts. **a**, Comparison of snuABE editing efficiency based on tagRNA length at the human *AAVS1* locus. **b**, Indel frequency and purity of the edited target adenine (A) derived from **a**. **c**, Editing efficiency comparison based on the base opposite the target adenine at the *AAVS1* and *FANCF* loci. The symbol (–) denotes a bulge. **d**, Effective targeting range of snuABE across 13 distinct sites within the *AAVS1* locus. **e**,**f**, Editing efficiency across all three-base motifs (NAN) at 32 endogenous loci. Sequence logos were generated using the ‘seqLogo’ package in R, based on normalized base editing efficiency rather than nucleotide frequency. **g**, Total base conversion rates of ABE8e and snuABE3.1 at six off-target sites identified by Cas-OFFinder. **h**, On-target vs. off-target editing activities of ABE8e and snuABE3.1 as measured by an orthogonal R-loop assay, based on the pooled data from three independent biological replicates, shown in **Supplementary Fig. 5b** for each target and off-target site. **i**, Comparison of A-to-I editing effects on seven endogenous mRNA sites using full-length human ADAR1, human ADAR2, snuABE1.0, and snuABE3.1. **j**, Whole-RNA sequencing analysis of A-to-I editing ratios for ABE8e, H840A, snuABE1.0, and snuABE3.1, each performed in a single replicate. The cell values in the heat map represent mean values; bar graphs show mean ± s.d. with error bars (*n* = 3 independent biological replicates). All experiments were performed in HEK293T cells.

ADAR is known to exhibit sequence preferences for RNA editing, favoring a 5’ U and 3’ G adjacent to the target adenine^31^. Therefore, we similarly examined the sequence context preferences of snuABE3.1. Using 32 target sites with varied 5’ and 3’ flanking sequences at adenines positioned between 12 and 15, we found that snuABE3.1 generally preferred A or T at the 5’ position and C or G at the 3’ position relative to the target adenine (**Fig. 3c**). Across these arbitrarily selected targets in HEK293T cells, the median editing efficiency was 2.1% (**Fig. 3f**), suggesting that snuABE3.1 is highly target-dependent. We also observed that the broader surrounding sequence context affects snuABE3.1 activity. For example, when editing an AAVS1-derived target containing a non-preferred motif (5’-GAC-3’), the editing efficiency was 17.3%, slightly lower but comparable to that of the preferred motif (5’-AAC-3’), which yielded 20.8% editing (**Supplementary Fig. 4**). However, altering the flanking distal sequences while retaining the preferred 5’-AAC-3’ led to a significant reduction in efficiency to 6.9%. These results indicate that while snuABE3.1 exhibits a clear motif preference, its editing efficiency is also significantly influenced by the broader sequence context.

### Determination of off-target effects on genome and transcriptome

Conventional ABEs have been reported to induce both gRNA-dependent^7^ and gRNA-independent off-target edits in the genome^9,10^, as well as gRNA-independent RNA deamination events^11–13,32^. Similarly, we investigated both tagRNA-dependent and tagRNA-independent off-target effects of snuABE3.1 on the genome and transcriptome. To assess tagRNA-dependent off-target activity, we tested snuABE3.1 at the six most likely off-target sites for *AAVS1*, which include sequences with up to two mismatches or one bulge. While ABE8e showed substantial off-target editing at all six sites, no detectable off-target DNA editing was observed with snuABE3.1 (**Fig. 3g** and **Supplementary Fig. 5a**). To evaluate tagRNA-independent genomic deamination, we employed the orthogonal R-loop assay ^33^. This assay involves co-expression of BEs with a catalytically inactive *Staphylococcus aureus* Cas9 (dSaCas9), which forms artificial R-loop structures, thereby revealing potential off-target deaminase activity independent of gRNA targeting. Notably, while ABE8e induced strong off-target editing at all dSaCas9 binding sites, snuABE3.1 showed no detectable off-target editing at any of the tested sites (**Supplementary Fig. 5b, 5c**), despite displaying comparable on-target editing efficiency at *AAVS1* (**Fig. 3h**).

RNA A-to-I editing, mediated by ADARs, is a frequent and functionally important event in human cells, playing a role in post-transcriptional gene regulation^16^. Given this, we hypothesized that snuABE3.1 might induce gRNA-independent RNA deamination. To evaluate this, we first analyzed six well-characterized endogenous ADAR hotspot sites in human mRNAs^34^. Although the editing levels were lower than those typically observed with native hADARs, snuABE3.1 still induced detectable A-to-I conversions at several of these sites (**Fig. 3i** and **Supplementary Fig. 3a**). To further assess transcriptome-wide RNA off-target activity, we performed whole-transcriptome RNA sequencing in cells expressing snuABE1.0 (containing hADAR2dd), snuABE3.1 (containing PhADARdd), ABE8e (positive control), and nCas9(H840A) (negative control). The results revealed that snuABE3.1 exhibited global A-to-I RNA editing frequencies higher than snuABE1.0 but comparable to ABE8e (**Fig. 3j**), indicating that snuABE3.1 has strong RNA deamination activity.

### Enhancing editing efficiency of snuABE through AI-based directed evolution

Although snuABE3.1 demonstrated usable levels of editing efficiency, its overall activity remained lower than that of conventional ABEs, and its sequence motif preference posed limitations. To address these issues, we focused on engineering the PhADARdd of snuABE3.1. Structural prediction of PhADARdd bound to a DNA:RNA hybrid using AlphaFold 3^35^ revealed a prominent N-terminal flap (**Fig. 4a**). We hypothesized that while this flap might be important for the full-length PhADAR protein, it may not be essential for the catalytic PhADARdd. Therefore, we generated a truncated version of PhADARdd lacking the N-terminal 10 amino acids, which resulted in enhanced editing activity (**Fig. 4b**). This improved variant was named snuABE4.0.

**Fig. 4.**
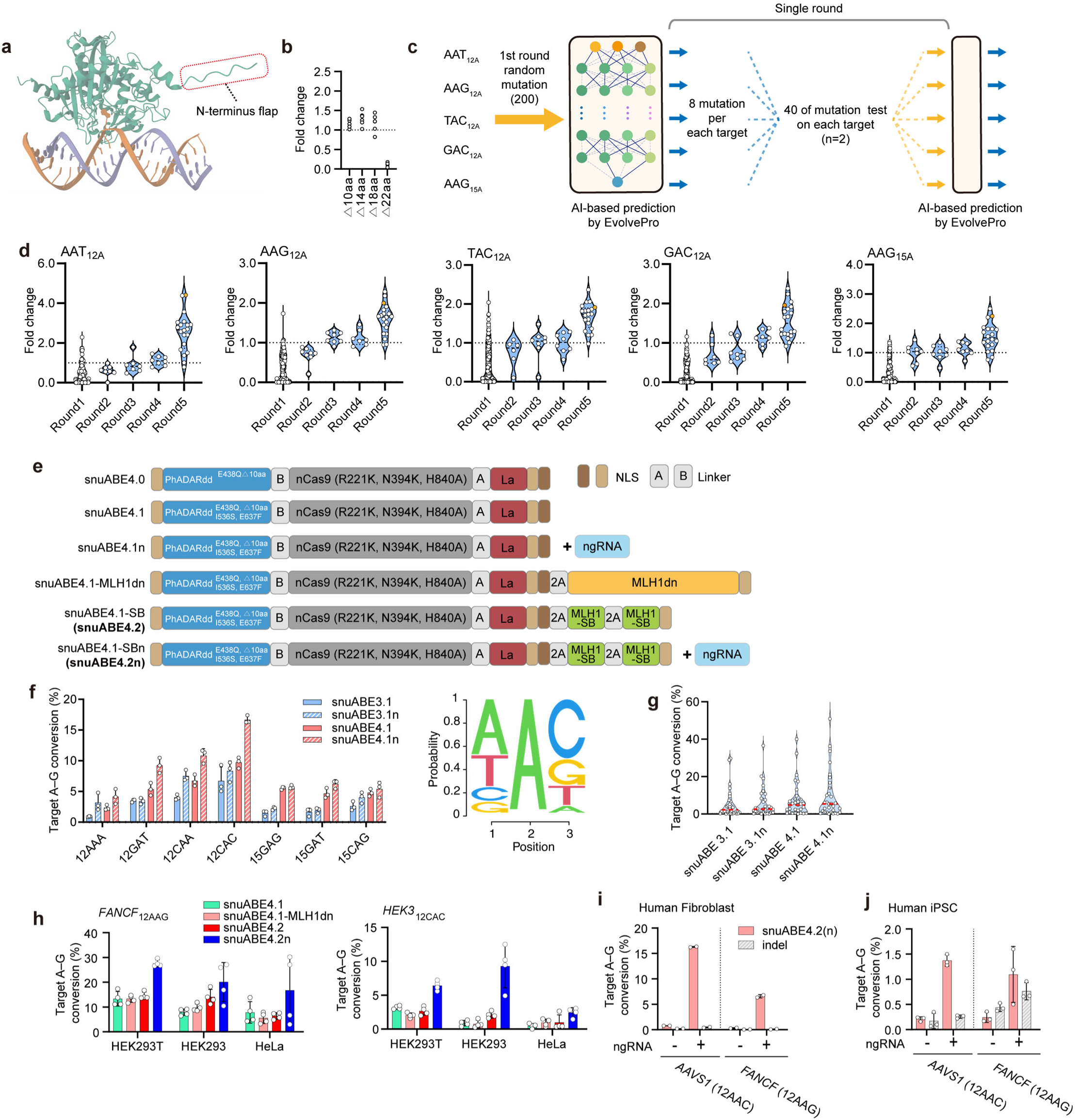
Engineering and optimization of snuABE variants via AI-assisted evolution. **a**, Predicted structure of the *P. humanus* ADAR deaminase domain bound to an DNA:RNA hybrid, modeled using AlphaFold3. DNA is shown in orange, RNA in blue. **b**, Editing efficiencies of truncated N-terminal flap variants of snuABE3.1. Each point represents a single replicate performed on a distinct target. **c**, Schematic of the directed evolution strategy using EvolvePro. Each point indicates the mean of two independent biological replicates per target. **d**, Fold-change in editing efficiency from predicted mutations identified by EvolvePro, tested across five endogenous targets. **e**, Schematic representations of snuABE4.0, snuABE4.1, snuABE4.1n, snuABE4.2, and snuABE4.2n, incorporating an additional nicking gRNA and MLH1-SB fusion. **f**,**g**, Motif-wide editing efficiencies of snuABE3.1, snuABE3.1n, snuABE4.2, and snuABE4.2n. Each data point shown in **g** individually represents the mean of each 35 targets presented in **Supplementary Fig. 6a,b**. **h**, Editing efficiency comparisons of these tools at the *FANCF* and *HEK3* loci across multiple cell lines (HEK293T, HEK293, and HeLa). **i**,**j**, A-to-G editing efficiencies of snuABE4.2(n) at the *AAVS1* and *FANCF* loci in human fibroblasts and induced pluripotent stem cells (iPSCs). The bar graphs show the mean ± s.d. with error bars; experiments shown in **f**,**j** were performed with n = 3 independent biological replicates, **h** with n = 4, and **i** with n = 2. Unless otherwise indicated, experiments were performed in HEK293T cells.

EvolvePro was used to further improve editing efficiency and reduce sequence bias in snuABE4.0. We first selected five endogenous genomic sites with diverse sequence motifs and target A positions (**Fig. 3e**): 5’-AAT-3’, 5’-AAG-3’, 5’-TAC-3’, 5’-GAC-3’ at position 12 (with editing efficiencies of 0.36%, 10.2%, 3.9%, and 1.02%, respectively), and 5’-AAG-3’ at position 15 (1.2% editing efficiency). In the first round, we generated 187 snuABE variants containing each sinlge random mutation in PhADARdd, and evaluated their performance into HEK293T cells. For each target, the top eight mutations were selected based on predicted editing gains by EvolvePro, resulting in 40 unique variants. These were experimentally tested across all five target sites, and the resulting data were used to iteratively retrain EvolvePro to improve prediction accuracy in subsequent rounds (**Fig. 4c**). Through successive rounds of evolution, we observed a stepwise improvement in editing efficiency. In the final (fifth) round, we utilized EvolvePro’s combinatorial mutation prediction module to identify the top 20 multi-mutant variants, each containing 2–5 mutations predicted to synergistically enhance activity. Experimental validation of these multi-mutants revealed a substantial gain in editing efficiency, with one variant harboring three mutations (E438Q, I536S, E637F) showing the highest activity. This optimized version was designated snuABE4.1 (**Fig. 4d**, **4e**). Notably, snuABE4.1 exhibited a significantly reduced motif bias, showing robust editing even at previously disfavored sequence contexts, such as targets with a 5’-G **(Fig. 4f)**. Additionally, overall editing efficiency improved markedly, with a median editing rate of 4.8% **(Fig. 4g)**, compared to 2.1% observed with snuABE3.1.

### MLH1-SB and additional nick further enhance editing efficiency of snuABE

The snuABE system utilizes nCas9(H840A), which introduces a nick on the same DNA strand as the target adenine. This configuration may limit editing efficiency during the repair process. Therefore, similar to the PE system (e.g., PE2 and PE3), we hypothesized that introducing an additional nick on the opposite DNA strand could enhance editing efficiency (**Fig. 4e**)^36^. To test this, we co-delivered nicking guide RNAs (ngRNAs) along with either snuABE3.1 or snuABE4.1, generating the modified versions snuABE3.1n and snuABE4.1n, respectively. These systems were tested across 32 target sites with various sequence motifs. The addition of the opposite-strand nick substantially improved editing outcomes, with snuABE4.1n achieving a median editing efficiency of 5.4% (**Fig. 4g**). Furthermore, motif dependency was alleviated in snuABE4.1n (**Fig. 4f**), supporting the effectiveness of the additional nick in enhancing snuABE performance.

We also considered the DNA repair pathways involved in snuABE activity. While conventional BEs primarily engage the base excision repair (BER) pathway, we posited that snuABE may also invoke mismatch repair (MMR) due to mismatches introduced on the nicked strand by nCas9(H840A), which could be targeted and corrected by the MMR system. To test this hypothesis, we co-expressed snuABE4.1 with either a dominant-negative MLH1 (MLH1dn) or a MLH1 small binder (MLH1-SB), both of which act as MMR inhibitors^25,37^. These configurations were termed snuABE4.1-MLH1dn and snuABE4.1-SB (also referred to as snuABE4.2), and were tested at the *FANCF* and *HEK3* loci. In MMR-deficient HEK293T cells ^38^, neither MMR inhibitor increased editing efficiency. However, in MMR-proficient HEK293 and HeLa cells, both inhibitors significantly enhanced editing efficiencies (**Fig. 4h**), indicating that suppression of MMR is beneficial for snuABE performance. Further improvement was achieved by combining ngRNAs with snuABE4.2, generating snuABE4.2n. This configuration led to a robust increase in editing activity across all tested loci (**Fig. 4h**). To evaluate its potential in clinically relevant contexts, we assessed snuABE4.2n activity in human fibroblasts and induced pluripotent stem cells (iPSCs) at the *AAVS1* and *FANCF* loci. Notably, editing efficiencies were also enhanced upon ngRNA addition, reaching up to 16.3% at the *AAVS1* locus in fibroblasts and 1.4% in iPSCs (**Fig. 4i**, **4j**).

### Precise adenine correction in genetic disease models by snuABE

To evaluate the therapeutic potential of snuABE, we first established a HEK293T cell model of Von Hippel-Lindau (VHL) syndrome (**Fig. 5a**) ^39^. To correct the target adenine mutation, three gRNAs were designed: one targeting a 5’-NG-3’ protospacer adjacent motif (PAM) and two targeting 5’-NGG-3’ PAMs. While conventional ABEs exhibited high levels of bystander editing within the editing window, snuABE4.1n achieved bystander-free, pinpoint editing with an efficiency of up to 63.6% (**Fig. 5b**). These results demonstrate that snuABE can target previously inaccessible genomic sites where conventional ABEs are limited by bystander effects, broadening the scope for therapeutic genome editing.

**Fig. 5.**
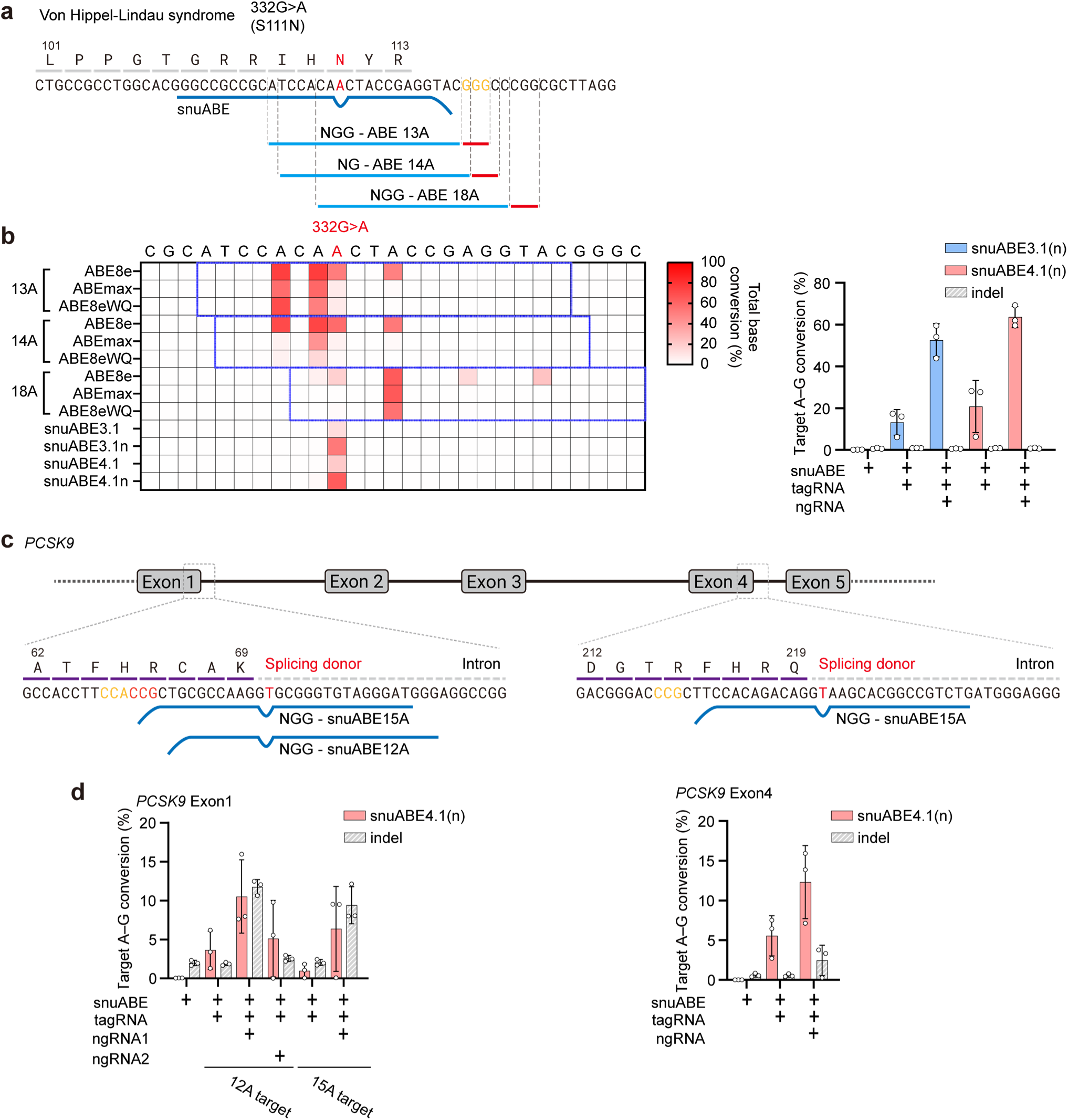
Application of snuABE variants for therapeutic A-to-G correction at disease-relevant genomic loci. **a**,**b**, Comparison of editing outcomes between conventional ABEs and snuABEs in a VHL syndrome-mimicking disease model using HEK293T cell lines. **c**,**d** Evaluation of snuABE-mediated A-to-G editing at the splicing acceptor region of the *PCSK9* gene in HeLa cells. The cell values in the heat map represent the mean, while the bar graphs show the mean ± s.d. with error bars (*n* = 3 independent biological replicates).

Low-density lipoprotein (LDL)-related disorders pose a major global health burden, and genome editing of *PCSK9* has emerged as a promising strategy to reduce LDL levels by disrupting *PCSK9* expression^40^. However, minimizing off-target effects is critical for clinical application. To this end, we investigated the suitability of snuABE, which is characterized by extremely low DNA off-target activity, for precise and safe *PCSK9* editing in HeLa cells. Splice donor sites located downstream of exons 1 and 4 in *PCSK9* were targeted using snuABE4.2, both with and without additional ngRNAs (**Fig. 5c**). Robust A-to-G conversions were achieved, reaching up to 10.5% at the exon 1 site and 12.3% at the exon 4 site (**Fig. 5d**), supporting the clinical applicability of snuABE for treating LDL-associated disorders.

## Discussion

Although conventional ABEs are widely used due to their ease of application and high editing efficiency, they are significantly limited by the frequent occurrence of bystander editing, which constrains their therapeutic utility. Various strategies have been proposed to address this limitation, including TadA engineering^14,28,41^, sgRNA modification^42^, motif-preferred ABEs^43^, and the use of accessory proteins^44^. However, these approaches have shown limited success in achieving precise single-base editing. In this study, we developed snuABE to achieve single-nucleotide resolution adenine conversion by inducing editing specifically at mismatched adenines formed in DNA:RNA hybrids. This design significantly reduced bystander editing. Furthermore, snuABE exhibited markedly lower DNA off-target activity, likely due to the requirement for perfect complementarity between the tagRNA and the target sequence, similar to the mechanism used by prime editors. These features position snuABE as a promising and safer BE for therapeutic applications. The recent clinical success of ABE-based therapy for carbamoyl-phosphate synthetase 1 (CPS1) deficiency highlights the translational potential of BEs and underscores the need for improved precision and safety^45^.

To optimize snuABE up to version 4.2, we adopted strategies from PE development, due to their similar architecture involving nCas9(H840A) and 3’-extended gRNAs. The additional mutations and NLSs used in PEmax were incorporated into snuABE2.1 ^25^. Likewise, introducing an additional nick on the opposite strand (as in the PE3 strategy) and protecting the 3’-end of tagRNA with La protein (as in PE7) contributed to the creation of the snuABE3 variants ^25,36^. Moreover, we applied the AI-based EvolvePro algorithm to engineer the PhADARdd deaminase, generating snuABE4 variants in a manner analogous to the PE6 strategy^22,46^. MMR suppression through MLH1dn or MLH1-SB further enhanced editing efficiency, paralleling the approaches used in PE4 and PE-SB, and led to the generation of snuABE4.2^25,37^.

Despite these improvements, snuABE4.2 still faces several limitations: (i) lower editing efficiency compared to conventional ABEs such as ABE8e, (ii) residual sequence preference around the target adenine, and (iii) considerable off-target editing on RNA transcripts. While RNA off-target effects are transient, these issues highlight the need for further refinement. Strategies such as genome-wide CRISPR screening to identify novel cofactors or advanced ADAR engineering, potentially via AI-guided discovery of orthologs, may offer promising paths forward. In addition, it would be beneficial to further develop snuABE to induce cytosine editing without bystander effects.

Nonetheless, snuABE provides a compelling alternative to conventional ABEs, offering high specificity and minimal DNA off-target activity. While prime editors are highly versatile and capable of inducing various types of edits (insertions, deletions, substitutions), their clinical utility is often hindered by the complexity of pegRNA design and typically lower editing efficiency. In contrast, snuABE relies on a relatively simple tagRNA design, requiring only optimization of the 3’ extension length (typically 24–32 nucleotides). In a disease-relevant model of VHL syndrome, snuABE achieved bystander-free editing with efficiencies comparable to those of ABE8e. Collectively, these results underscore the potential of snuABE to therapeutically target genomic loci that remain inaccessible using current ABE technologies.

## Acknowledgements

The authors thank Dr. Francisco X. Mora (EssayReview) for English-language editing. Most of the sequencing data analysis was conducted using the computing server at the Genomic Medicine Institute Research Service Center. This research was supported by grants from the National Research Foundation of Korea (NRF) (No. 2021M3A9H3015389, No. RS-2024-00451880, No. RS-2024-00455559, and SRC-NRF2022R1A5A102641311) awarded to S.B. Additional support was provided by the Korean Fund for Regenerative Medicine (KFRM) (No. RS-2024-00332601), a grant from the Ministry of Food and Drug Safety (No. 25202MFDS003) in 2025, and the SNUH Lee Kun-hee Child Cancer & Rare Disease Project (No. 25B-001-0700), also awarded to S.B.

## Author contributions

H.W.I. and S.B. conceived the project. H.W.I. and C.-J.J. developed the bioinformatics algorithms. H.W.I., B.-D.J., Y.L., Y.E.O., and Y.-W.K. performed the cell experiments. H.W.I. and H.U. carried out the protein engineering. S.B. supervised the project. H.W.I. and S.B. wrote the manuscript with input from all authors.

## Additional information

Supplementary Information accompanying this paper is available at http://

## Materials and Methods

### Molecular cloning of tagRNAs, agRNA, sgRNA, and ngRNA

tagRNAs, sgRNAs, and ngRNAs used for plasmid transfection were cloned into the pU6-pegRNA-GG-acceptor plasmid (Addgene, 132777). anchor-guide RNAs (agRNAs) used with ABEx1–4^30^ were cloned into the pU6-tevopreq1-GG-acceptor plasmid (Addgene, 174038). The vectors were digested with BsaI-HFv2 (New England Biolabs, R3733L). Spacers and extensions were synthesized as oligonucleotides (Macrogen), phosphorylated using T4 Polynucleotide Kinase (enzynomics, M005S), and annealed (37 °C for 30 min; 95 °C for 2 min; cooling to 25 °C at a 0.1 °C/s for 5 minutes). The annealed oligos were ligated into the digested vectors using T4 DNA ligase (enzynomics, M001S) and transformed into *E. coli* DH5α competent cells prepared with Mix & Go (Zymo Research, T3001).

### Cloning of snuABE constructs

All snuABE constructs were derived from ABE8e (Addgene, 138489). Two rounds of site-directed mutagenesis were performed to introduce the A10D reversion and H840A mutation into Cas9. The resulting plasmid was digested with NotI and BglII (NEB, R0189S, R0144S), and the ADAR deaminase domains were inserted using Gibson Assembly® Master Mix (NEB, E2611L). Human ADAR1 and ADAR2 were obtained from pDY1173_ADAR1p150 (Addgene, 193192) and pmGFP-ADAR2 (Addgene, 117929), respectively. The *Drosophila* ADAR sequence was obtained from the HyperADAR control plasmid (Addgene, 166969). ADAR deaminase domains from other species were synthesized (Twist Bioscience). The PEmax domain used to construct snuABE2.1 was derived from pCMV-PEmax (Addgene, 174820). The La protein CDS used in snuABE3.1 was synthesized from HEK293T cDNA by PCR. Total RNA was extracted from HEK293T cells (ATCC, CRL-11268) using Trizol (Invitrogen™, 15596026), and cDNA was synthesized using ReverTra Ace (Toyobo, FSK-101F). The CDS was then amplified using KOD plus Neo (Toyobo, TOKOD-401).

### Mammalian cell culture conditions

HEK293T (ATCC, CRL-11268), HEK293 (ATCC, CRL-1573), HeLa (ATCC, CLL-2), and U2OS (Korean Cell Line Bank, KCLB, Seoul, Republic of Korea) cells were cultured in DMEM (Welgene, LM001-05) supplemented with 10% FBS (Welgene, PK004) and 1% antibiotics (Welgene, LS203-1) at 37 °C in a 5% CO_2_ incubator. For passaging, cells were washed with DPBS (Welgene, LB001-01), detached using 0.25% Trypsin-EDTA (Welgene, LS015), and resuspended in fresh media before being transferred to new plates. Human fibroblasts were maintained in DMEM supplemented with 10% FBS, 1% antibiotics, and 1× GlutaMAX™ Supplement (Gibco, 35050061) at 37 °C with 5% CO_2_. Cells were cultured in T75 flasks (SPL, 70075) with 20 mL of medium and passaged using TrypLE™ Express (Gibco, 12605010) after washing with DPBS. hiPSCs (CMC-hiPSC-022, Catholic University of Korea) were cultured in Essential 8™ medium (ThermoScientific, A1517001) on 60-mm tissue culture dishes coated with Vitronectin (Gibco™, A14700). For passaging, cells were washed with DPBS and detached using ACCUTASE™ (Stem Cell Technologies, 07922). After detachment, cells were washed twice with DPBS and transferred to Vitronectin-coated plates in Essential 8™ medium supplemented with ROCK inhibitor (Y27632 dihydrochloride, MCE, HY-10583). hiPSCs were maintained at 37 °C with 5% CO_2_.

### Transfection and transduction protocols and genomic DNA extraction

For HEK293T, HEK293, HeLa, and U2OS cells, 0.25 × 10^5^ cells were seeded per well in 48-well plates (SPL, 30048) containing 250 µL of DMEM supplemented with 10% FBS and 1% antibiotics, one day prior to transfection. The following day, plasmid vectors were mixed and transfected using 0.5 µL of jetOPTIMUS (Polyplus, 101000006) per well, according to the manufacturer’s instructions. Typically, 375 ng of snuABE and 125 ng of tagRNA were used per transfection. If required, an additional 41.6 ng of ngRNA was included (541.6 ng in total). When co-expressing additional proteins such as La protein or MLH1dn, 100 ng of plasmid was added, bringing the total to 600 ng. After 36 hours, 200 µL of medium was replaced with fresh medium (DMEM, 10% FBS, 1% antibiotics). Cells were harvested 36 hours later using 0.25% Trypsin-EDTA (Welgene, LS015).

For transfection of human fibroblasts, electroporation was performed using the Neon® Transfection System (Invitrogen™) with the 10 μL Kit (Invitrogen™, MPK1096). Cells detached using TrypLE™ Express (Gibco™, 12605010) were resuspended in Neon™ R buffer at a density of 1 × 10^5^ cells per reaction and mixed with 1 μg of total plasmid DNA. Electroporation was carried out at 1600 V with a 20 ms pulse width and a single pulse. After 72 hours, cells were harvested using TrypLE™ Express.

For hiPSC transfection, 1 × 10^5^ cells were seeded into a 24-well plate. Plasmid DNA (same total amount as above) was diluted in 25 μL of Opti-MEM™ (Gibco™, 31985-070), then mixed with 25 μL of Opti-MEM™ containing 2 μL of Lipofectamine Stem (ThermoFisher, STEM00015). The resulting 50 μL mixture was transfected into the cells. At 72 hours post-transfection, cells were washed with DPBS and detached using ACCUTASE™ (Stem Cell Technologies, 07922).

Harvested cells were centrifuged at 1000 × g for 5 minutes to collect the pellet. Genomic DNA was extracted by adding 100 µL of extraction buffer (40 mM Tris, pH 8.9; 1% Tween-20; 0.2 mM EDTA; Proteinase K [20 mg/mL]; 0.2% Nonidet P-40). Samples were vortexed for 15 seconds and incubated in a thermal cycler at 60 °C for 15 minutes, followed by 98 °C for 5 minutes.

### Amplicon sequencing and data analysis

Genomic DNA was analyzed using an Illumina MiniSeq sequencer. Targeted deep sequencing was performed using a two-step PCR process. In the first step, 2 µL of extracted DNA was amplified using adaptor primers. In the second step, 1 µL of the initial PCR product was used as the template with index primers. PCR was conducted with KOD multi & Epi (Toyobo, KME-101) following the manufacturer’s instructions under the following cycling conditions: 98 °C for 2 minutes; 29 cycles of 98 °C for 10 seconds, 56 °C for 20 seconds, and 68 °C for 30 seconds; and a final extension at 68 °C for 1 minute. The final PCR products were purified using Expin™ PCR SV (GeneAll Biotechnology, 103-102) and sequenced using the MiniSeq Mid Output Kit (Illumina, FC-420-1004). Analysis of A-to-I editing efficiency at target sites was performed using the ABE_detector pipeline (https://github.com/imhw4157/ABE_detector), and total base conversion rates were calculated using the allsub_detector pipeline (https://github.com/imhw4157/allsub_detector).

### RNA sequencing and data analysis

The Neon® Transfection System (Invitrogen™) with the 100 μL Kit (Invitrogen™, MPK10096) was used for mRNA sequencing. HEK293T cells were detached as previously described, resuspended in DPBS, and washed by centrifugation at 500 g for 3 minutes. A total of 1 × 10^6^ HEK293T cells were resuspended in 100 µL of sucrose potassium phosphate buffered saline (119 mM KCl, 15.1 mM NaCl, 0.144 mM CaCl_2_, 5 mM MgCl_2_, 9.98 mM KH_2_PO_4_, 250 mM Sucrose, pH 7.4), mixed with 2 µg of vector, and subjected to electroporation (1600 V, 20 ms pulse width, 1 pulse). The transfected cells were transferred to 6-well plates (SPL, 30006) containing 2 mL of DMEM supplemented with 10% FBS and incubated at room temperature for 5 minutes in a clean bench, followed by incubation at 37 °C with 5% CO_2_ for 24 hours. After incubation, the media was removed entirely, and total RNA was extracted using 1 mL of TRIzol (Invitrogen™, 15596026) and 200 µL of Phenol:Chloroform:Isoamyl Alcohol (25:24:1, saturated with 10 mM Tris, pH 8.0, and 1 mM EDTA; Sigma-Aldrich, P2069-100ML).

For targeted RNA sequencing (**Fig. 3i,j**), cDNA was synthesized using ReverTra Ace (Toyobo, FSK-101F) according to the manufacturer’s protocol and sequenced using Illumina MiniSeq as previously described. For whole-RNA sequencing (**Fig. 3k**), total RNA was processed using the NEBNext® rRNA Depletion Kit v2 (Human/Mouse/Rat) and RNA Sample Purification Beads (NEB, E7405L), followed by library preparation using the NEBNext® Ultra™ II RNA Library Prep Kit with Sample Purification Beads (NEB, E7775S), according to the manufacturer’s instructions. Sequencing was outsourced to Novogene (NovaSeq X Plus), with 6 Gb of sequencing data generated per sample.

For RNA sequencing data analysis, FASTQ files were aligned to the GRCh38 reference genome (primary assembly from Ensembl) using HISAT2^47^ (https://github.com/DaehwanKimLab/hisat2). The resulting BAM files were sorted and indexed using SAMtools^48^ (https://www.htslib.org). A-to-I RNA editing events were analyzed using REDItools2^49^ (https://github.com/BioinfoUNIBA/ REDItools2) following the published protocol. To ensure data reliability, editing sites with a read depth of less than 50 were excluded using the -l (MIN_COLUMN_LENGTH) option in REDItools2.

### Generation of Lenti knock-in HEK293T cells

To generate HEK293T cells with an artificial motif knock-in, lentiviral transduction was employed. The lentiviral vector was constructed by digesting pAX198 (Addgene, 173042) with XbaI and XhoI (NEB, R0145S and R0146S), following the same cloning strategy used for tagRNAs, agRNAs, sgRNAs, and ngRNAs. Lenti-X™ 293T cells (Takara, 632180) were seeded at 6 × 10^6^ cells per 100 mm cell culture dish (SPL, 20100), and the medium was replaced with fresh DMEM containing 10% FBS the following day. Immediately after media replacement, a mixture consisting of 9.4 µg of the constructed lentiviral vector, 8.6 µg of psPAX2 (Addgene, 12260), 2.6 µg of MD2.G (Addgene, 12259), and 2.0 µg of pRSV-Rev (Addgene, 12253) was prepared and transfected using 65.7 µL of Transporter 5® Transfection Reagent (Polysciences, 26008-50), according to the manufacturer’s instructions. One day after transfection, the media was replaced with fresh DMEM containing 10% FBS. Two days later, the media containing the viral particles was collected, centrifuged at 500 g for 10 minutes to remove cellular debris, and filtered through a 0.45 µm filter (Pall, 4614). The virus was then concentrated using the Lenti-X™ Concentrator (Takara, 631232) following the manufacturer’s protocol. HEK293T cells were infected with the concentrated lentivirus at a multiplicity of infection (MOI) of 0.3. Two days post-infection, cells were selected with 3 µg/mL puromycin (InvivoGen, ant-pr-1) for three days.

### Directed evolution through EvolvePro

Directed evolution using the EvolvePro pipeline was conducted as previously described (https://github.com/mat10d/EvolvePro)^22^. Approximately 200 random single-point mutations were generated and tested in HEK293T cells across five independent target sites. The editing efficiencies of these variants were measured and used as training input for the initial round of EvolvePro predictions. For each target, the top eight mutations predicted to exhibit the highest activity were selected, resulting in a total of 40 variants (8 per target × 5 targets), which were then experimentally validated across all five sites in subsequent rounds. This iterative process was repeated over multiple rounds, with EvolvePro models updated using the most recent experimental data in a target-specific manner. In the final round, all mutation data were compiled, and the average fold-change in editing efficiency across the five targets was calculated. Mutations showing a fold-change greater than 1 were retained and used to generate all possible combinatorial variants containing up to four mutations. These multi-mutant variants were ranked using EvolvePro, and the top 20 predicted variants were selected for final experimental validation.

