## Supplementary Information for "Engineered ADARs enable single-nucleotide resolution DNA A-to-G editing without bystander effects"

**List of Supplementary Materials:**

Supplementary Figure 1 to 6.

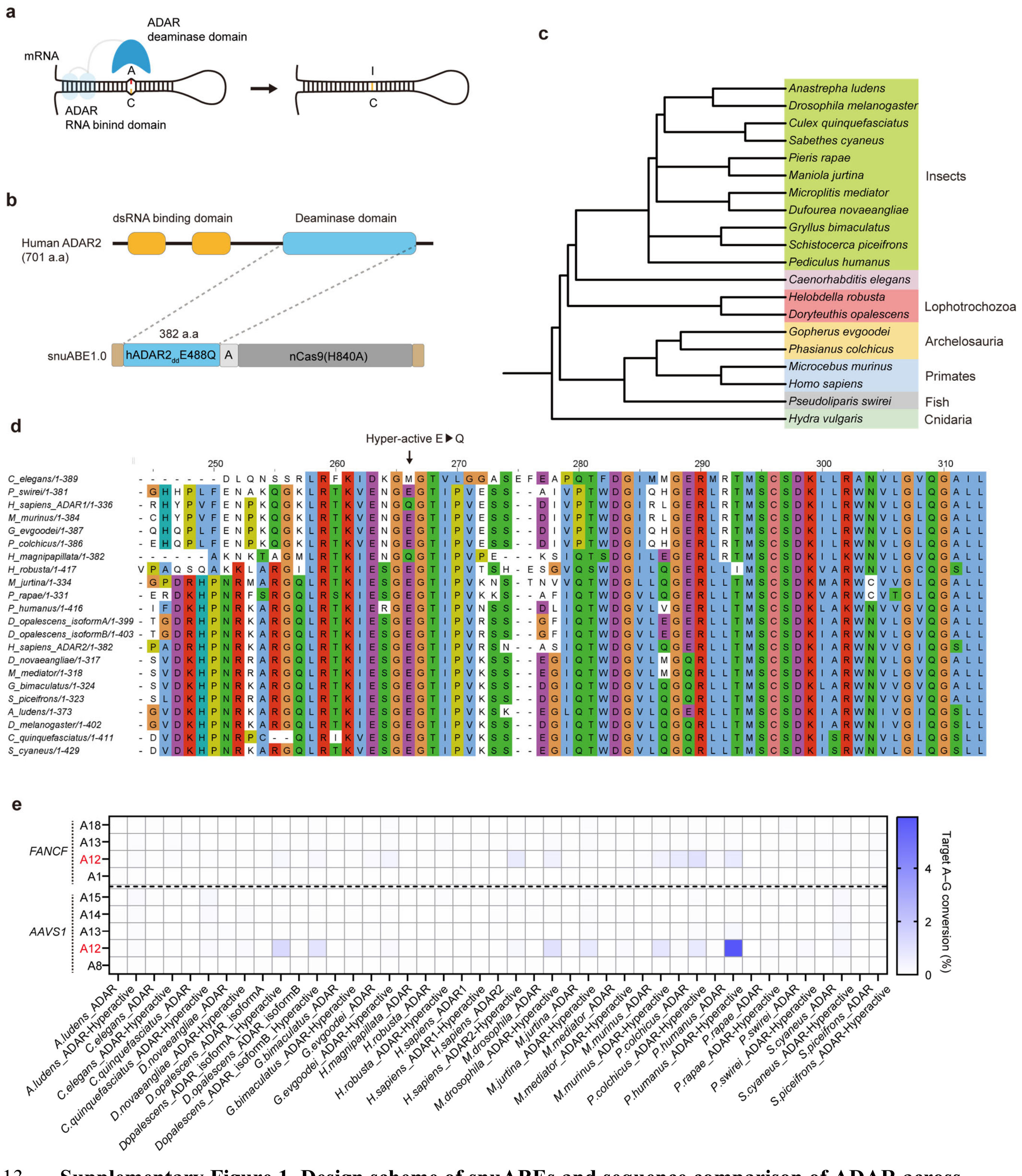


**Supplementary Figure 1. Design scheme of snuABEs and sequence comparison of ADAR across various species. a**, Schematic of the native ADAR protein functioning in mRNA. **b**, Schematic of full-length human ADAR2 protein and snuABE1.0. **c,** Phylogenetic tree of the species used in **Fig. 1d**. **d**, Alignment of amino acid sequences from the ADAR deaminase domains of the species in **Fig. 1d**. **e**, Editing efficiency comparison of the various ADAR proteins tested at the human *AAVS1* and *FANCF* loci. All experiments were conducted in HEK293T cells. The cell values in the heat map represent the mean ($n = 3$ independent biological replicates).

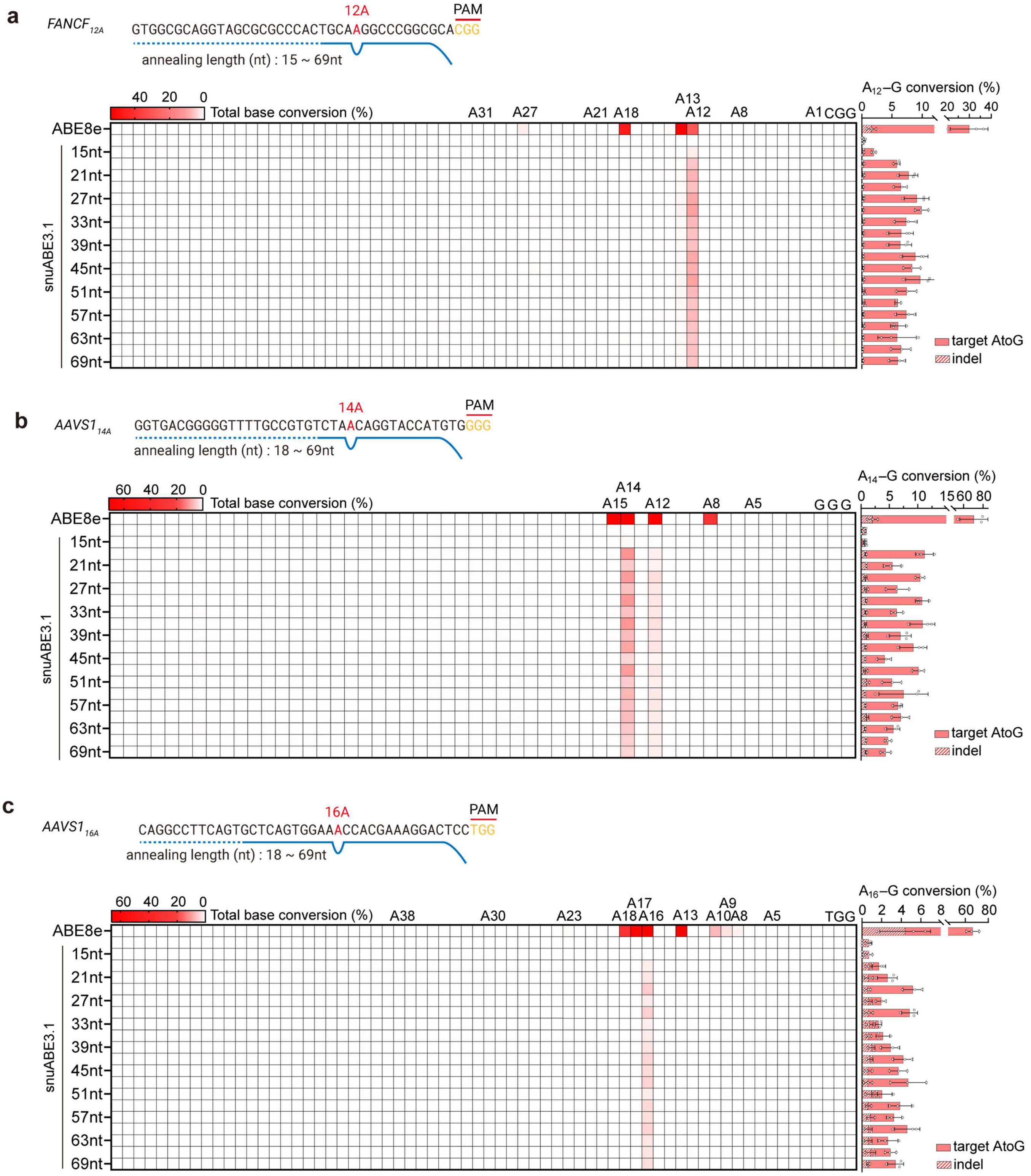


**Supplementary Figure 2. Efficiency comparison of snuABE3.1 across various targets based on tagRNA length. a-c**, Efficiency comparison of snuABE3.1 with varying tagRNA lengths across three independent targets. The cell values in the heat map represent the mean, while the bar graphs show the mean ± s.d. with error bars ($n = 3$ independent biological replicates). All experiments were conducted in HEK293T cells.

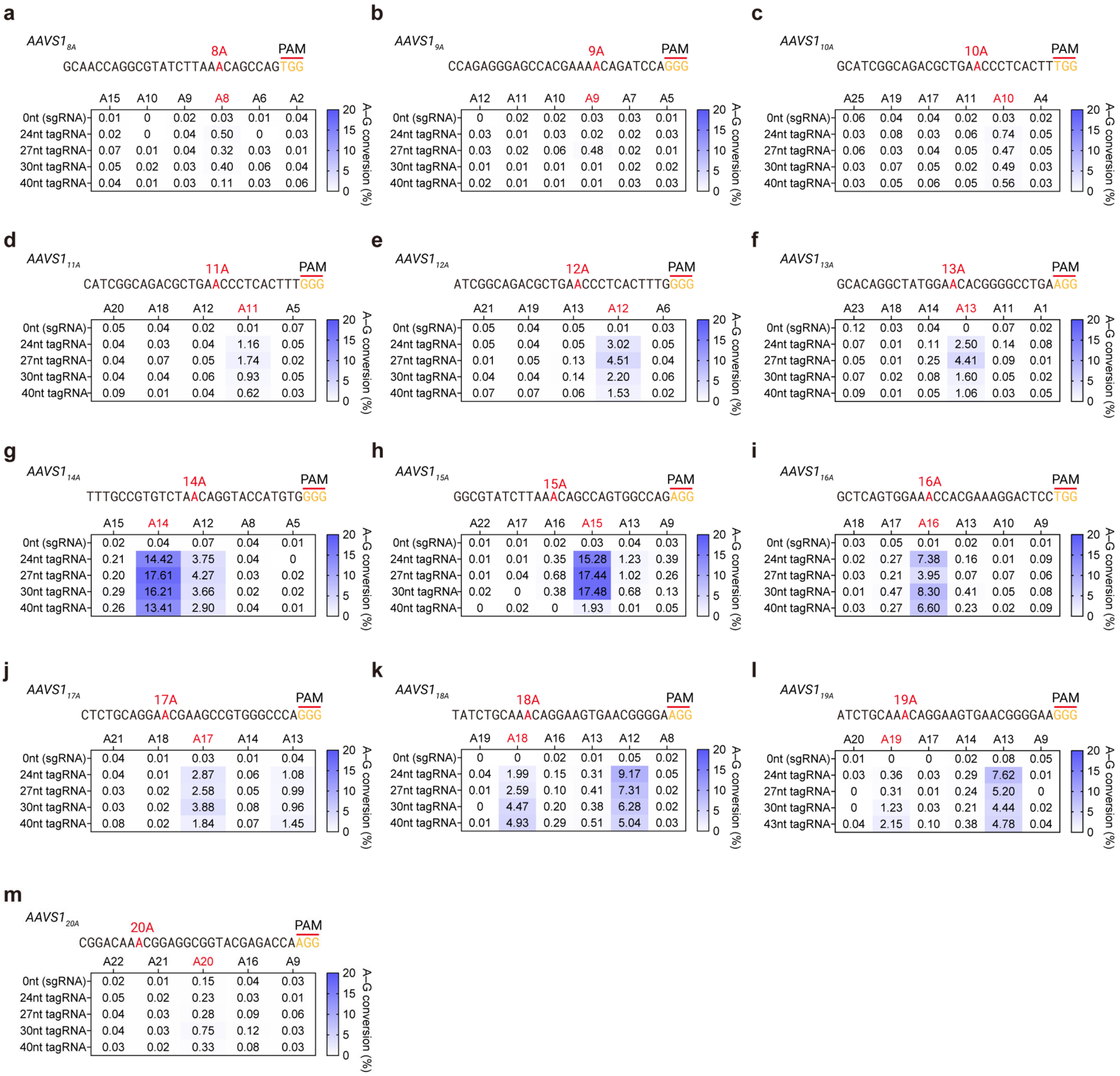


**Supplementary Figure 3. Efficiency comparison of snuABE3.1 at different target adenine positions. a-m**, Editing efficiency of snuABE3.1 at target positions A8 to A20 across thirteen independent sites. The cell values in the heat map represent the mean ($n = 3$ independent biological replicates). All experiments were conducted in HEK293T cells.

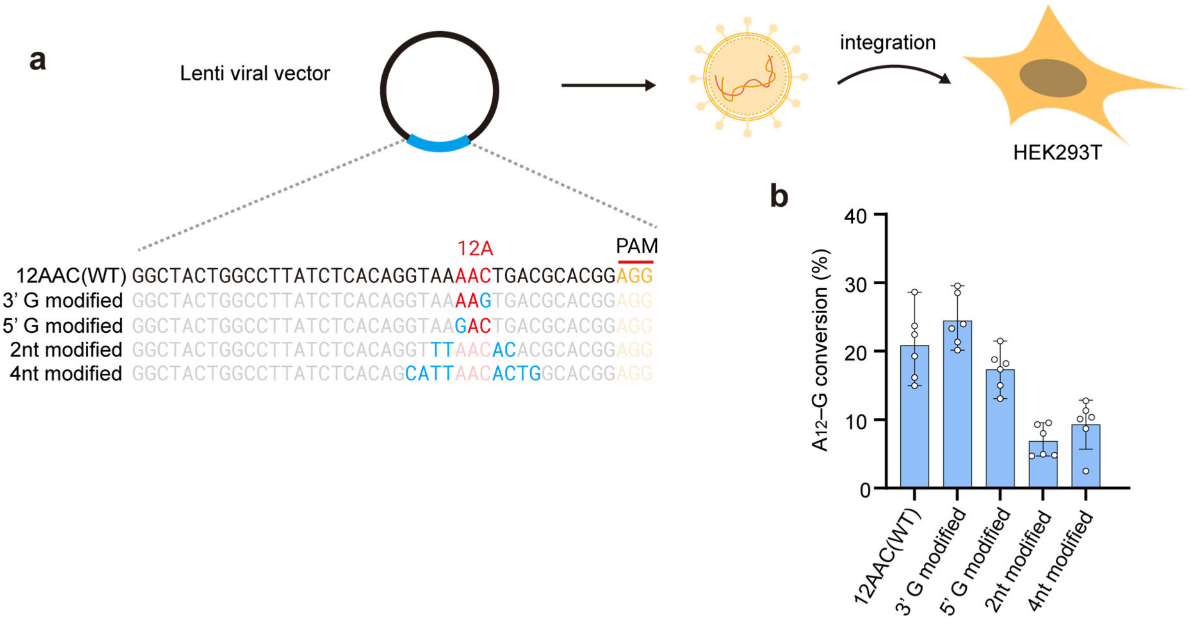


**Supplementary Figure 4. Efficiency comparison of snuABE3.1 on artificial targets with modified motifs. a**, Schematic showing the generation of artificial motifs. **b**, Editing efficiency of snuABE3.1 tested on the artificial motifs from a. Bar graphs show the mean ± s.d. with error bars. (*n* = 6 independent biological replicates). All experiments were conducted in HEK293T cells.

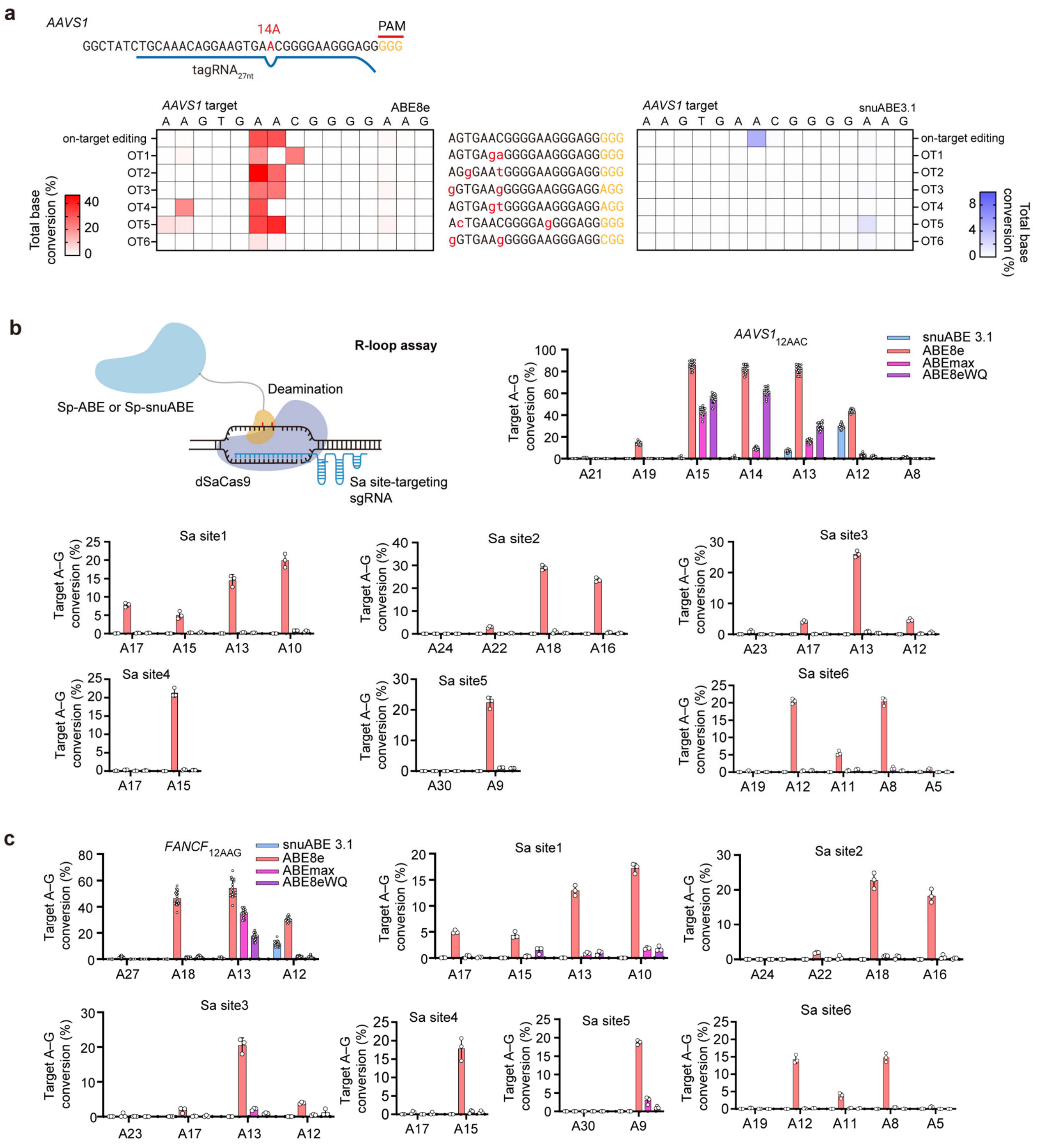


**Supplementary Figure 5. : Cas-dependent and -independent DNA off-target effects of snuABE3.1.** **a,** Comparison of total base conversion rates for ABE8e and snuABE3.1 at six Cas-OFFinder-predicted off-target sites. **b**,**c**, Orthogonal R-loop assay evaluating off-target A-to-G editing by ABEs at the *AAVS1* and *FANCF* loci, as well as six dSaCas9 binding sites. The cell values in the heat map represent the mean, while bar graphs show the mean ± s.d. with error bars (*n* = 3 independent biological replicates). All experiments were conducted in HEK293T cells.

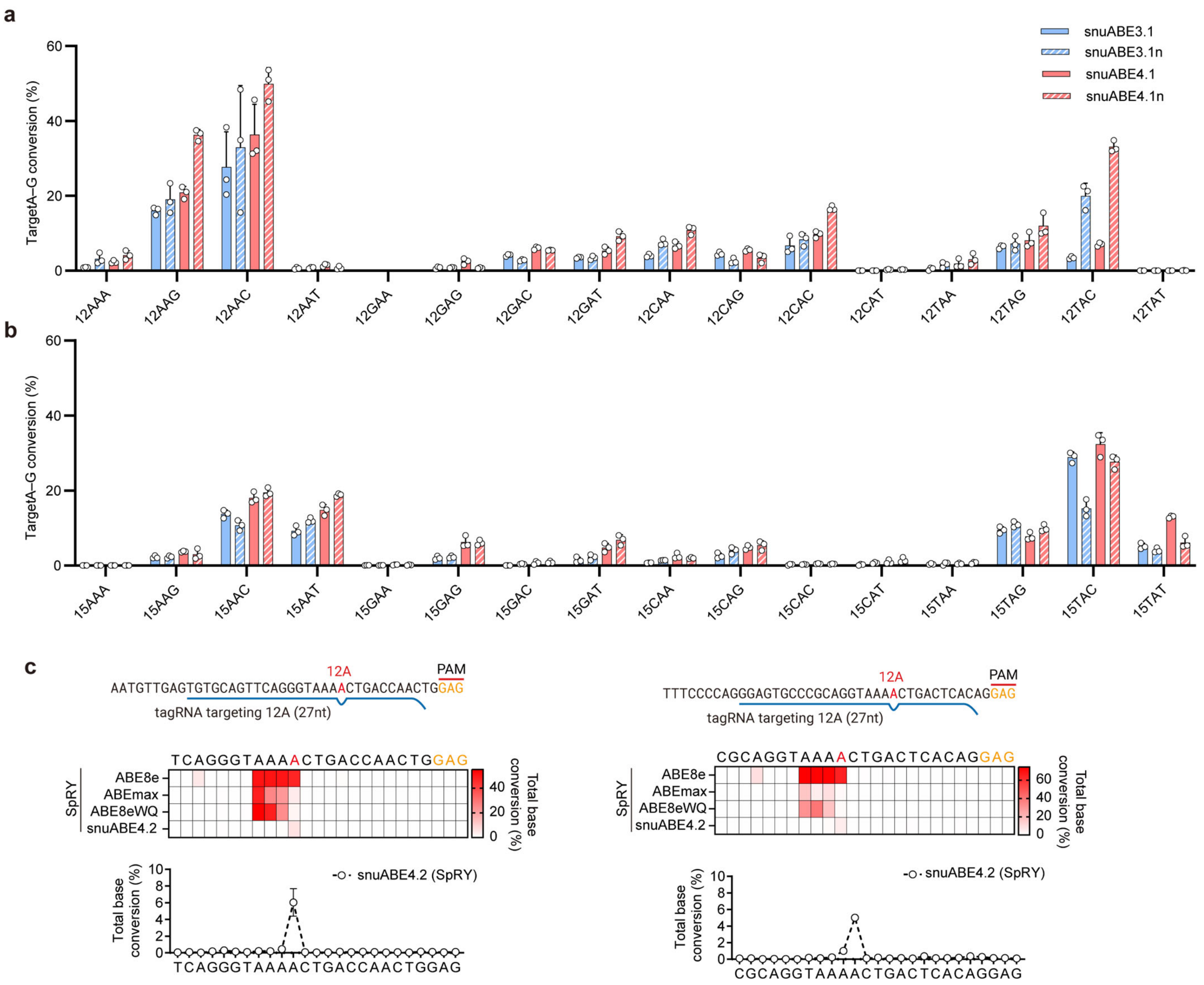


**Supplementary Figure 6. Motif-wide editing activity and PAM-less target recognition of snuABE(SpRY). a**,**b**, Editing rates of target adenines across all 16 possible NAN motifs at A12 **a** and A15 **b**, using snuABE3.1, snuABE3.1n, snuABE4.1, and snuABE4.1n. **c**, Editing windows of different ABE variants (ABE8e, ABEmax, ABE8eWQ) and snuABE4.2 combined with the SpRY system. The cell values in the heat map represent the mean, while bar graphs show the mean ± s.d. with error bars (*n* = 3 independent biological replicates). All experiments were conducted in HEK293T cells.
